# The Major Histocompatibility Complex modulates *Batrachochytrium dendrobatidis* and *Ranavirus* infections in three amphibian species

**DOI:** 10.1101/2023.04.14.536887

**Authors:** M Cortazar-Chinarro, A Richter-Boix, P Halvarsson, G Palomar, J Bosch

**Affiliations:** MEMEG/Department of Biology, Lund University, Sölvegatan 37, SE-22362 Lund, Sweden; Animal Ecology/ Department of Ecology and Genetics, Uppsala University, Norbyvägen 18D, SE- 75236 Uppsala, Sweden; Department of Earth Ocean and Atmospheric sciences/ Faculty of Science 2020 – 2207 Main Mall Vancouver, BC Canada V6T 1Z4; Department of Political and Social Science, Pompeu Fabra University, Ramon Trias Fargas 25-27, 08005 Barcelona, Spain; Parasitology/Department of Biomedical Sciences and Veterinary Public Health, Swedish University of Agricultural Sciences, Ulls väg 26, P.O. Box 7036, SE-75007 Uppsala, Sweden; Department of Genetics, Physiology and Microbiology, Complutense University of Madrid, C/Jose Antonio Novais 12, Madrid 28040, Spain; Instituto Mixto de Investigacion en Biodiversidad, Universidad de Oviedo, Campus de Mieres. Edificio de Investigacion-Planta 5, c/ Gonzalo Guiterrez Quiŕs s/n, 33600 Mieres, Asturias

**Keywords:** amphibian, co-infections dynamics, Chytrid fungus, viruses, infection status

## Abstract

Genetic variation of immune genes is an important component of genetic diversity. Major histocompatibility complex (MHC) genes have been put forward as a model for studying how genetic diversity is maintained and geographically distributed in wild populations. Pathogen-mediated selection processes (i.e., heterozygosity advantage, rare-allele advantage or fluctuating selection) and demography are believed to generate and maintain the extreme diversity of MHC genes observed. However, establishing the relative importance of the different proposed mechanisms has proved extremely difficult, but heterozygote advantage is expected to be more detectable when multiple pathogens are considered simultaneously. Here, we test whether MHC diversity in three amphibian species (*Ichthyosaura alpestris, Pleurodeles waltl,* and *Pelophylax perezi*) is driven by pathogen-mediated selection. We examined the relationship between the individual MHC class II exon variability with individual infection status (infected or not), infection intensity, and co-infection of two main amphibian pathogens: *Batrachochytrium dendrobatidis* (*Bd*) and *Ranavirus* sp. (*Rv*). We found higher MHC class II exon 2 allelic diversity in *I.alpestris* and *P. perezi* than in *P.waltl* but no significant differences in allele frequencies between infection groups. We also observed significant differences in *Bd* infection intensity between *Bd* infected individuals and co-infected individuals depending on the number of MHC loci that an individual carries. For *I. alpestris*, we show stronger evidence for MHC associations with infection intensity and status when individuals carry specific alleles and supertypes. Our results suggest that studying the association between MHC genes and single and co-infected individuals might provide new insights into host-parasite evolution and a better understanding of evolutionary mechanisms driven by MHC diversity.

## INTRODUCTION

Host-pathogen coevolution is known to drive the maintenance of genetic variation in immune genes, with consequences for important evolutionary processes including the evolution of virulence, kin recognition (Sommer, 2005), and sexual selection (Grieves, Gloor, Bernards, & MacDougall-Shackleton, 2019; van Oosterhout, 2009; Winternitz et al., 2013). The major histocompatibility complex (MHC) is an example of genetic polymorphism thought to be maintained by such processes: subjected to strong diversifying and balancing selection, which leads to the maintenance of high diversity in natural populations (Spurgin & Richardson, 2010). Despite its wide ecological and evolutionary importance, the processes driving evolution at the MHC are not fully understood (Radwan, Babik, Kaufman, Lenz, & Winternitz, 2020). There are several mutually non-exclusive evolutionary mechanisms that may contribute to the maintenance of MHC diversity after pathogen-imposed selection in populations (Jan Ejsmond, Radwan, & Wilson, 2014).

The heterozygosity advantage hypothesis assumes that individuals that possess a diverse MHC perform best, a greater number of alleles increases the spectrum of antigens recognized providing a selective advantage in pathogen recognition (Doherty & Zinkernagel, 1975; Penn, Damjanovich, & Potts, 2002). Consistently with the heterozygote mechanisms, several studies reported a positive linear relationship between MHC diversity and different fitness traits (Thoss, Ilmonen, Musolf, & Penn, 2011; Whittingham, FreemanLGallant, Taff, & Dunn, 2015), and negatively associated with the prevalence or intensity of infection by different parasites (Radwan et al., 2012; Slade, Sarquis-Adamson, Gloor, Lachance, & MacDougall-Shackleton, 2017). However, an excess of heterozygotes may also be compatible with rare-allele advantage or negative frequency-dependent selection, as heterozygotes may be selected for because they carry rare or novel MHC variants (i.e. resistant alleles) rather than because they are heterozygous *per se* (Spurgin & Richardson, 2010). Further theorical work have suggested that frequency-dependent selection resulting from Red Queen dynamics may be an important process (Ejsmond & Radwan, 2015) but it has only been demonstrated recently in the context of natural MHC diversity in experimental studies and long-term monitoring studies (Migalska et al., 2022; Phillips et al., 2018). Indeed, the number of studies reporting associations between MHC variation and spatial differences in parasite communities are leading to increase (O’Reilly, Paterson, Mitchell, & Gardner, 2023; Osborne, Pilger, Lusk, & Turner, 2017; Radwan et al., 2020).

An opposite mechanism is the hypothesis of immunogenetic optimality (Nowak, Tarczy-Hornoch, & Austyn, 1992), where the wider recognition of pathogens by a higher heterozygosity is counteracted by the increased risk of autoimmune diseases and the reduction of functional T-cells at high MHC diversity at the individual level (Migalska et al., 2022; Woelfing, Traulsen, Milinski, & Boehm, 2009). This hypothesis results in an optimal number of genes at intermediate individual MHC diversity what revealed advantages in certain individuals (Kalbe et al., 2009; Wegner, Kalbe, Kurtz, Reusch, & Milinski, 2003). Several studies found support for the MHC optimality hypothesis in mammals, birds, reptiles, and fishes (Bonneaud, Mazuc, Chastel, Westerdahl, & Sorci, 2004; Hablützel et al., 2014; Kloch, Babik, Bajer, Siński, & Radwan, 2010; Madsen & Ujvari, 2006), especially in the three-spined stickleback, in which individuals with an intermediate number of MHC loci suffered least from simultaneous infections with multiple pathogens in wild populations (Wegner et al., 2003). However, there is also a considerable number of studies that did not detect benefits of such intermediate MHC diversity (Radwan et al., 2012; Sepil, Lachish, & Sheldon, 2013), raising the question whether the benefits from intermediate diversity depend on the species studied and/or the intensity of parasitic infection (Milinski, 2006). For instance, directional selection occurs when a specific allelic lineage, that confers resistance to a common pathogen, increases in frequency over successive generations; there is some evidence that at least sometimes strong selection from pathogens may in fact reduce MHC polymorphism (de Groot et al., 2002; Savage & Zamudio, 2016; Teacher, Garner, & Nichols, 2009). Finally, if the pathogen regime faced by a population fluctuates spatial-temporally, the intensity of directional selection at MHC genes will also fluctuate, leading to the fluctuating selection hypothesis. This hypothesis proposes that spatial and temporal heterogeneity in the type of parasites and pathogens may maintain diversity (Hedrick, 2002; Spurgin & Richardson, 2010).

Understanding the extent to which immunogenetics variation modulates disease outcomes in wild populations is critical to understand their ability to adapt and persist, especially when it comes to emerging diseases. Novel emerging infectious diseases continue appearing all over the world in a great variety of taxa, being identified as an increasing threat to wildlife conservation (Cunningham, Daszak, & Wood, 2017; Tompkins, Carver, Jones, Krkošek, & Skerratt, 2015; Woolhouse, 2008). Large-scale epidemics have affected vertebrates such as birds, mammals (Sacristán et al., 2021), and reptiles (Lorch et al., 2016). However, the severity and extent of the impact of the emerging diseases chytridiomycosis and ranavirosis in amphibians are unprecedented in the history of vertebrates (Fisher & Garner, 2020; Miller, Gray, & Storfer, 2011; Scheele et al., 2019; Skerratt et al., 2007). Chytridiomycosis is caused by the fungal pathogens *Batrachochytrium dendrobatidis* (*Bd*) and the more recently discovered *Batrachochytrium salamandrivorans* (*Bsal*) (Martel et al., 2014), whereas ranavirosis is originated by the Irridoviridae viral lineage *Ranavirus* (Brunner, Storfer, Gray, & Hoverman, 2015). Recent evidence indicates that co-infection by both pathogens is common (Herczeg, Ujszegi, Kásler, Holly, & Hettyey, 2021; Olori et al., 2018), but little information exists on within-host interactions between these agents and how populations adapt to both of them. While some studies suggest that co-infection has a positive effect on each other (Stutz et al., 2018; Whitfield et al., 2013), others have not confirmed this interaction (Warne, LaBumbard, LaGrange, Vredenburg, & Catenazzi, 2016) or even found negative effect (Bosch, Monsalve-Carcaño, Price, & Bielby, 2020).

To date, the interplay of chytridiomycosis and host immune MHC diversity has been charaterizated in several amphibian species. The majority of the Mstudies focused on investigating the relation between MHC class II diversity (e.g. nucleotide and amino acid sequence polymorphism) and infection (Kosh et al. 2019), finding evidence of positive selection in MHC alleles (Bataille et al., 2015; May, Zeisset, & Beebee, 2011; Savage & Zamudio, 2016) or heterozygosity advantage (Savage & Zamudio, 2011). Extreme polymorphism found in MHC class II genes in amphibians is belived to evolve in response to pathogen diversity (Savage & Zamudio, 2011; Sommer, 2005). While many studies have focused on MHC class II relationship to *Bd*, the evidence of the relationship between MHC II and *Ranavirus* susceptibility was lacking until recently (Savage *et al*, 2019) and studies on co-infections dynamics and how multi-host parasite affect the immunological response are still scarce, even if co-infection with multiple pathogens is a real world rule (Hoarau, Mavingui, & Lebarbenchon, 2020).

Here, we investigated whether and how the selective mechanisms associated to the two most devastating pathogens of amphibians (*Bd* and *Rv*) could drive the genetic diversity of MHC class II genes in three amphibian species. We specifically studied both pathogen prevalences and infection intensities in relation to MHC diversity and heterozygosity. We also tested whether specific MHC configurations are associated with higher or lower susceptibility in an attempt to disentangle the different pathogen-mediated selection mechanisms, even if it is difficult (Spurgin & Richardson, 2010).

## METHODS

### Sample collection, DNA extraction, and infection detection

We sampled 69 Alpine newts (*Ichthyosaura alpestris*), 57 Iberian ribbed newt (*Pleurodeles waltl*), and 51 Iberian green frog (*Pelophylax perezi*) in two locations of the Iberian Peninsula: Picos de Europa and Zamora (Figure 1; Supplementary Figure S1). Both locations are considered a focal source of single and co-infected sites where all infection categories are represented: 1) *Bd* infected, 2) *Rv* infected, 3) *Bd*+*Rv* infected and 4) non-infected (See Table 1). These locations are optimal to investigate co-infection dynamic trends due to the massive amphibian loss occurred there lately (J. Bosch pers. com.). We collected tissue samples (tail-clips) from adult individuals for *Bd* and *Rv* diagnostic tests. Lesions consistent with clinical signs of ranavirosis (erythema or hemorrhage) on the sampled individuals were noted.

**Figure 1.**
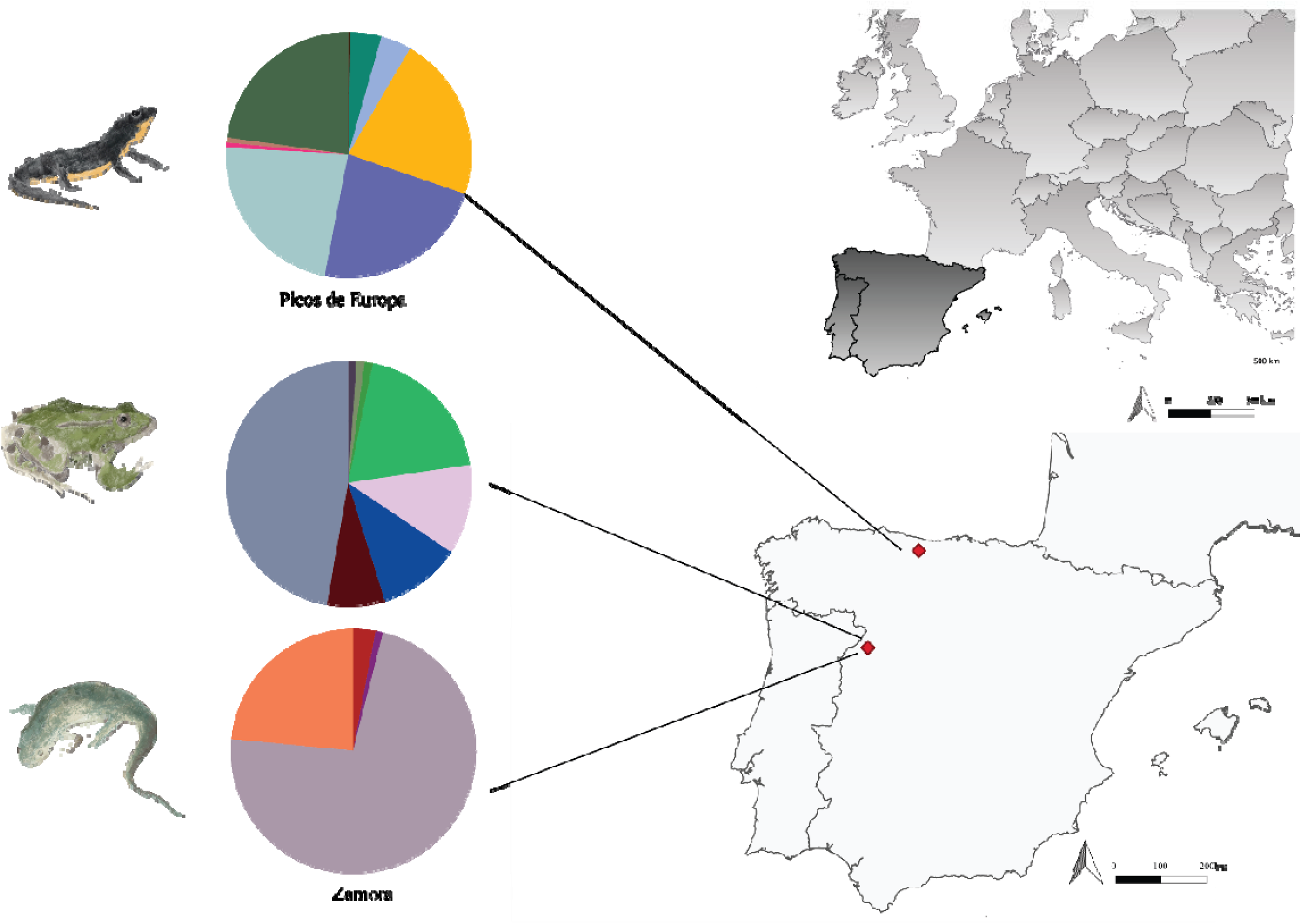
Allele frequency distribution of MHC class II exon 2 in *I. alpestris*, *P. perezi* and *P. waltl* in two locations (Picos the Europa and Zamora). Colour coding scheme for the alleles is given in the (Supplementary Figure S1). Amphibians illustrations were created by A.Cortazar for this specific study.

**Table 1.**
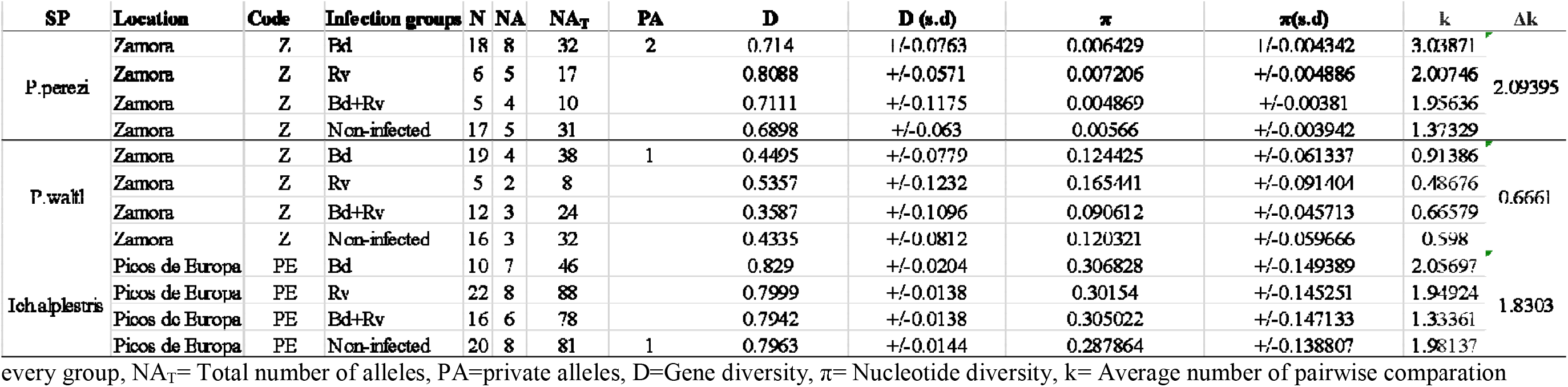
Genetic variation at the MHC class II exon 2 locus in Zamora and Picos de Europa locations. N=number of individuals, NA= number of alleles in every group, NA_T_= Total number of alleles, PA=private alleles, D=Gene diversity, π= Nucleotide diversity, k= Average number of pairwise comparation

Genomic DNA was extracted from 177 individuals (N=69 *I. alpestris,* N=57 *P. waltl,* and N=51 *P. perezi*) by using DNeasy Blood and Tissue Kit (Qiagen, Hilden, Germany). Purity and concentration of DNA were determined using NanoDrop® 2000 spectrophotometer and Qubit®3.0 fluorometer Quantitation Kit (Invitrogen™). Two qPCRs, one for *Bd* and one for *Rv,* were performed on a MyGo Mini PCR machine at the MNCN-CSIC lab following Boyle et al. (2004) and Leung et al. (2017) respectively . Samples were run in duplicate and when they showed signs of inhibition (i.e. non-sigmoidal amplification) they were diluted to 1:100 (*Bd*) or 1:10 (*Rv*) and re-analyzed. Samples were considered positive when both replicates revealed infection loadsL>L0.1 (*Bd*) orL>L3 (*Rv*), and the amplification curves have a sigmoidal shape. If not, the sample was re-run and considered positive only with another positive result recorded. Pathogen load was measured as the mean of the two replicates from the same individual.

### MHC II exon 2 gene

#### Single locus amplification and preparation for sequencing

Different sets of MHC class II exon 2 primers were used for each of the species. For the *I.alpestris* and *P.waltl*, we used the primers TrMHCII1F (5’-TCTCTCCGCAGTGGACTTCGTG-3’) and TrMHCII5R (5’- CTCACGCYTCCGSTGCTCCATG -3’) designed by Babik et al. (2008). We amplified a partial fragment of the second exon (corresponding to the β −2 domain), 239 bp in *I.alpestris* and 355 bp in *P.waltl*. From those 355 bp fragment, we retained 270 bp for a valid allele assignation and 190 bp for MHC supertyping analyses of *P.waltl*. For *P.perezi*, we used ForN (5’- CCGCAGATGATTTC-3’) developed by Mulder et al. (2017) and RevM (5’- TCCGGTACTCACATTTCAGGTCT-3’; Mulder pers.comm). We amplified 252 bp of the complete MHC class II exon 2. The primers used in this study were not used previously for MHC class II exon 2 amplification in P*.waltl* and P*.perezi*. Therefore, PCR products were visualized and isolated from agarose gels (1.5%). The resulting product was sequenced for DNA-sequence and fragment length verification. All PCR products were standard Sanger-sequenced at Macrogen Europe (Netherlands). Both forward and reverse primers were modified for Illumina MiSeq sequencing with an individual 8 bp barcode and a sequence of three N (to facilitate cluster identification). Each sample was marked with an individual combination of a forward and a reverse barcode for identification. PCR reactions were conducted in a total volume of 20 μl containing 1 μl of genomic DNA, 2 μl of 10X Dream taq buffer (Thermo scientific lab), 0.4 μl of 2 mM of each dNTP, 0.5 μl of each 10 μM primer (TrMHCII1F/TrMHCII5R and ForN/RevM, respectively), 1.5 μl of Bovine Serum Albumine (BSA; 5 mg/ml), and 0.25 μl of Dream taq DNA polymerase (5 U/μl, Thermo scientific lab) in deionized water. Thermocycling was performed on an ABI 2720 thermocycler (Applied Biosystems®). An initial denaturation step of 3 min at 95 °C was followed by 30 s at 95 °C, annealing for 30 s at 70 °C (*I.alpestris* and *P.waltl*) and 62 °C (P*.perezi*) for a total of 30 cycles and ending with an extension step at 72 °C for 1 min. All amplifications were carried out using filter tips in separate (pre- and post-PCR) rooms, and negative controls were included in all amplifications to avoid contaminations following the exact protocol of Cortazar-Chinarro et al. (2018). PCR products were run and visualized on a 1.5% agarose gel using gelgreen (BIOTIUM). To reduce the number of samples for subsequent purification, 3–9 PCR products with similar concentrations were pooled based on estimations from the gel image. These sample pools were run on 1.5% agarose gel, the target band was excised from the gel and extracted using the MinElute Gel Extraction Kit (Qiagen® Sollentuna, Sweden). The concentration of each sample pool was measured with Quant-iT PicoGreen dsDNA assay kit (Invitrogen Life Technologies, Stockholm, Sweden) on a fluorescence microplate reader (Ultra 384; Tecan Group Ltd., Männedorf, Switzerland). The final amplicon pooling was done according to the measured concentrations and consisted of equimolecular amounts of each sample. A total of six amplicon pools were generated per run, and libraries were prepared using the Illumina Truseq DNA PCR-Free Sample preparation kit (Illumina Inc., San Diego, CA). Six pools were combined into a Miseq run (PE250) and sequencing of two Miseq runs was carried out at the National Genomic Infrastructure (NGI), the SNP&SEQ Technology Platform hosted at SciLifeLab in Uppsala (Sweden).

#### Miseq data analyses

Sequencing data were extracted from the raw data and paired-end reads were combined into single forward reads using FLASH (Magoč & Salzberg, 2011), each of the 12 amplicon pools was analyzed independently. In total, 12 fastq files were generated and transformed into fasta (multifasta) files using Avalanche NextGen package (DNA Baser Sequence Assembler v4 (2013), Heracle BioSoft). We used the AmpliSAS software (Sebastian et al. 2016) for demultiplexing, clustering, and filtering the amplicon sequencing data corresponding to *I.alpestris* and P*.perezi*. Due to the complexity of the Miseq amplicon data for P*.waltl* (i.e. variable amplicon size), we used DADA2 software for demultiplexing, clustering, and filtering (Callahan et al., 2016).Some of the sequences met the minimal overlap criteria and merged properly while others were lost in the process. To solve this issue, we used both “justConcatenated=TRUE” and “verbose=TRUE” option for retrieving most of good quality reads that differed in length. . The decision to use only the initial 190 base pairs was made due to the low quality control of the reverse read, as well as the insufficient overlap between the forward and reverse read sequences. Subsequently, the 190 base pairs were used for allele calling. Samples were run in duplicates or triplicates to ensure a high replication rate validation of complex samples. We discarded samples with <L300 reads from the analysis for quality reasons. In addition, we used the DOC method (Lighten et.al 2014) implemented in AmpliSAS, not assuming any specific number of loci to identify and estimate the number of alleles (Ai) per individual. This procedure is based on the break point in sequencing coverage between alleles within each individual and avoids choosing a subjective threshold to separate true alleles from artefacts. Alleles are sorted top-down by coverage, followed by the calculation of the coverage break point (DOC statistic) around each allele. The allele with the highest DOC value is assumed to be the last true allele (see (Lighten, Van Oosterhout, Paterson, McMullan, & Bentzen, 2014).

All valid alleles were imported to MEGAX (Kumar, Stecher, Li, Knyaz, & Tamura, 2018) to create an alignment, translated into aminoacids, and to identify sequences containing stop codons and consecutively discarded. All sequences were extensively compared to existing sequences for the same locus in related species: smooth newt (*Lissotriton montandoni*; GenBank: JN565330.1), northern crested newt (*Triturus cristatus*; GenBank: FJ448027.1), tiger salamander (*Ambystoma tigrinum*; GenBank: DQ071906.1), and moor frog (*Rana arvalis*; (Cortázar-Chinarro et al., 2017). Valid MHC exon 2 alleles were named following the nomenclature suggested by Klein 1975: abbreviation of the species name followed by gene*numeration, e.g., Ich_alp_DAB*01.

### Data Analyses

#### MHC II exon 2 Diversity

Relative allele frequencies were estimated for each MHC class II exon 2 allele by species and infection group (*Bd* infected, *Rv* infected, *Bd*+*Rv* infected and non-infected; (See Figure S1 and S2) using ARLEQUIN v. 3.5 (Excoffier & Lischer, 2010). To test differences in allele frequencies between infection groups at each location, we run an Analyses of Molecular Variance (AMOVA) test in ARLEQUIN v. 3.5 (Excoffier & Lischer, 2010). Allele frequency plots were created in R using the “ggplot2” package (Wickham, 2011). Standard diversity indices (H_E,_ expected heterozygosity; π, nucleotide diversity; k, average number of pairwise comparison nucleotide differences) were calculated per infection groups by species to assess genetic diversity in the MHC II exon 2 in ARLEQUIN v. 3.5 (Excoffier & Lischer, 2010); Table 1). The number of loci per species was defined by counting the total number of alleles per species and illustrated via a phylogenetic tree based on the Neighbor joining method with bootstrapping (1000 replicates, Supplementary Figure S3) implemented in MEGA7 (Kumar et al. 2016). We reconstructed an unrooted phylogenetic network in the software SplitTree4, using the neighbor net method (Allman, Kubatko, & Rhodes, 2017), Supplementary Figure S4) to illustrate the phylogenetic relationships among the alleles from each amphibian species. We include a *Rana arvalis* sequence from (Cortázar-Chinarro et al., 2017), as an outgroup.

#### MHC II exon 2 Supertyping

To collapse MHC alleles into functional supertypes, we extracted the MHC II exon the amino acid positions of PBRs (Peptide Binding Region) of the Human Leukocyte Antigen (HLA; MHC) based on a previous study lead by (Papadopoulos, Bondinas, & Moustakas, 2008) (See the amino acid alignment; Supplementary Figure S5). Then, we characterized each site based on five physio-chemical descriptor variables: z1 (hydrophobicity), z2 (steric bulk), z3 (polarity), z4, and z5 (electronic effects). A hierarchical clustering tree was constructed based on the PBRs of MHC class II exon 2 Human and the z-descriptors in R (version 4.0.5). The optimal number of clusters were decided based on divergence between the branches in the phylogenetic tree. Alleles within clusters were collapsed into a single Supertype (See Figure 2). Supertype allele frequency plots were created in Excel. For *I. alpestris*, the supertypes were grouped into specific supertype_haplotypes taking into consideration that different supertypes were present at the same time (See Figure 2 and Suplementary Figure S6). The Supertype_haplotype consist on four digits and every digit indicate the number of supertypes from 1 to 4 that is the maximum number of supertypes present in *Ich.alpestris* for a specific haplotype designation (e.g., Ich_alp*2_1_0_0). MHC supertype_haplotype frequencies were visualized using the ggplot2 package (Wickham, 2011) (Suplementary Figure S6).

**Figure 2.**
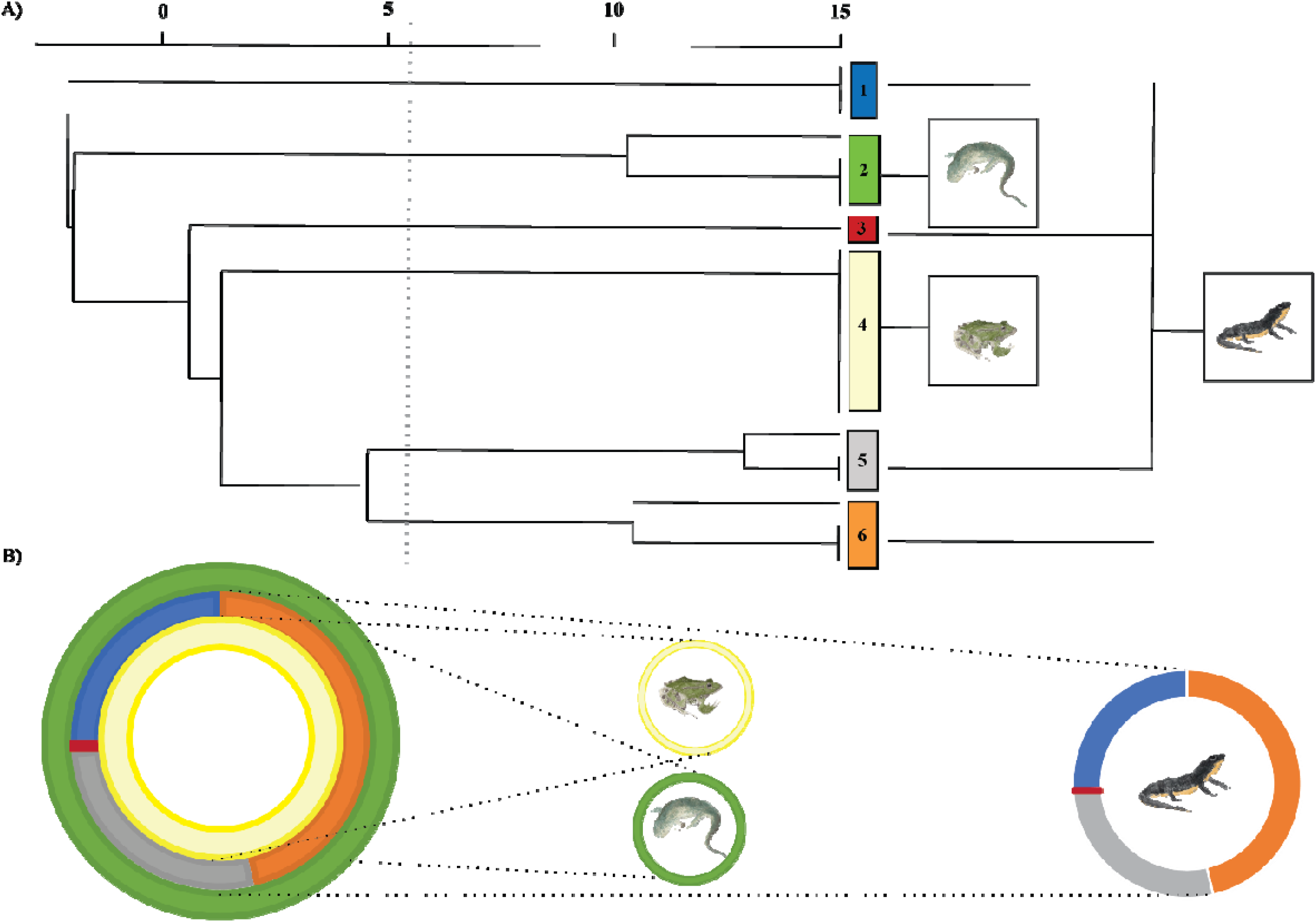
Phylogenetic dendogram reconstruction. A) Every colour squared represents a functional MHC class II exon 2 supertype. The grey line represents the threshold line stablished to cluster the aminoacid MHC class II sequences in functional supertypes. B) Pie charts represent the supertype frequency in the three species of the study: *I. alpestris*, *P. perezi* and *P. waltl*.). Amphibians illustrations were created by A.Cortazar for this specific study.

#### MHC II exon 2 and infection

A preliminary visual inspection of general patterns of infection by species and location was performed using the ggplot2 R package (Wickham, 2011); See Figure Supplementary Figure S7). Infection load was standardized to be able to compare results between *Bd* and *Rv* infection. In order to assess the importance of infection between species, we built a generalized linear mixed model (GLMM) based on the data distribution. We used *Bd*/*Rv* infection load as the response variable and species as the independent variable. For an accurate adjustment of the method, we implement Zero inflated Gamma distribution with a logit link. Additionally, we built a generalized linear mixed model (GLMM) to assess whether the infection intensity (*Bd* load, *Rv* load) was associated with the number of MHC II exon 2 loci and the infection group (Supplementary Figure S8). We used *Bd* load/*Rv* load as the response variable with the number of loci, Infection load and its interaction as independent variables. The model was controlled by “Species’, used as random factor,’ because different species differed in number of MHC loci . For a better adjustment of the model, we implement the Zero inflated Gamma, Gamma method with a log and inverse link, respectively (See Figure 3). The models were run by using the R CRAN package (glmmTMB) developed by Magnusson et al. (2017). We assessed the best model based on: 1) overdispersion, 2) homogeneity of variance and 3) the normality of residuals by using Dharma package (Hartig & Hartig, 2017). For obtaining the significant values of the fix effect in GLMM, we used Wald’s test.

**Figure 3.**
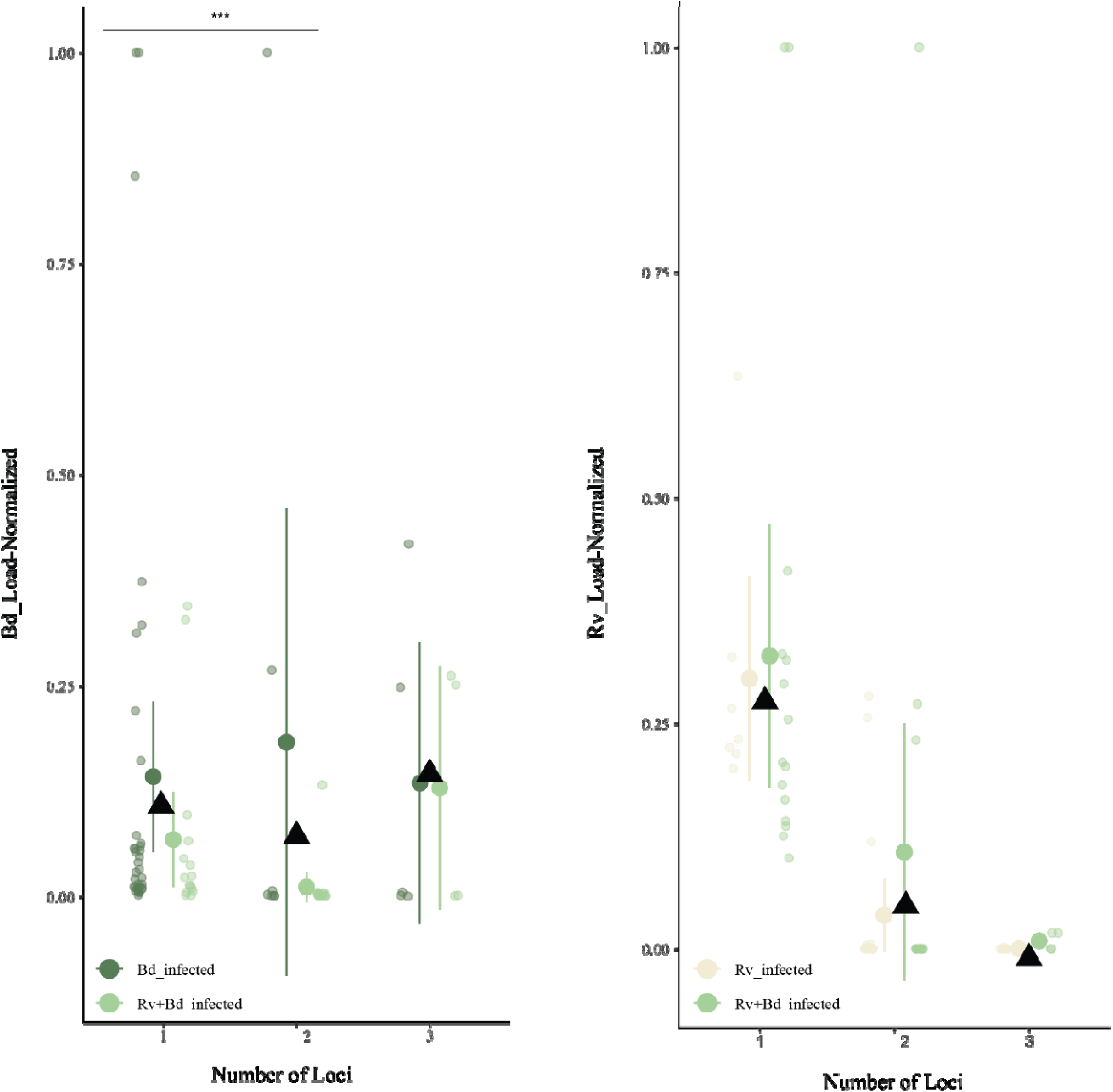
Infection loads normalized measures of Genome Equivalents (“GE”) for A) Bd and B) Rv infection in relation with the number of MHC class II exon 2 loci found in the study (a max of three), considering single infection (Bd or Rv) and co-infection (Rv+Bd). Black triangles represent the average of infection load per Number of Loci. Significant interactions (p-value<0.001) are represented with a black (*).

#### The effect of alleles on infection

To investigate whether specific individual MHC configuration is related to the infection status in the wild, we built generalized linear models (GLM). We run the models separately for each of the species, as no alleles were shared between species. *P.walt* was excluded from the analyses because of the low number of alleles, it could generate false positive/negative relationships between infection status and alleles. First, we included Infection status (infected or not) as the response variable with the specific alleles (e.g. Ich_alp *10+ Ich_alp *06+ Ich_alp *11+ Ich_alp *07+ Ich_alp *04+ Ich_alp *05) as independent variables. Second, we include either *Bd* infection status or *Rv* infection status as the response variable and the specific alleles as independent variables. Co-infected individuals were assigned to both *Bd* and *Rv* infection, respectively. We used a binomial error distribution with the logit link function in order to assess whether there was a relationship between carrying a specific allele and co-infection. Coefficient plots representing the effect of specific alleles on infection were constructed using dotwhisker (Solt & Hu, 2015) and broom packages (Robinson, 2014) implemented in R (See Supplementary Figure S9).

#### The effect of supertypes

To investigate whether specific MHC supertypes were associated to infection in *I. alpestris, P. perezi,* and *P. waltl*, we run GLMM models, controlling by species (random factor) with glmmTMBR package developed by (Magnusson et al., 2017). We used infection status as the response variable and the supertypes as the independent variable. Then, we focused on *I. alpestris* for consequent analyses on haplotypes_supertypes, as we obtained significant differences between infection status and carrying a specific supertype in this species. We build GLM models to investigate further relationships between specific haplotype-supertypes and infection status/intensity. We represented the regression coefficient plots for a better visual perspective, showing the main effect of carrying a specific supertype and haplotype- supertype over infection. We used dotwhisker (Solt & Hu, 2015) and broom packages (Robinson, 2014) implemented in R (See Supplementary Figure S10 and Supplementary Figure S11). Finally, we illustrated descriptive visual plots for haplotype_supertype and infection group, using the ggplot2 package in R (Wickham, 2011); Supplementary Figure S12).

## RESULTS

### Miseq run summary

The total number of reads obtained varied from 2,100,296 to 3,575,602. We obtained an average of 5,675,898 reads ± 1,107,689 (s.d.) with intact primer barcode information from two separate runs (Table S1). The average number of reads per amplicon varied depending on the species: 2591.90 ±117.59 (s.d.), 29,003.58 ± 340.99 (s.d.) and 23,336.62 ± 328.35 (s.d.) in *P. waltl*, *P. perezi,* and *I. alpestris*, respectively. For MHC class II exon2, we amplified and sequenced a total of 177 individuals in duplicates (N=21 *I. alpestris*, N=114 *P. waltl,* and N=57 *P. perezi*; Table S1). We include replicated individuals for each of the two Miseq runs (32 to 100%; Table S1). Amplicons with <L300 reads and replicates not producing the same alleles in each sample were discarded. The variation in number of duplications were connected to the success of the primer used. We assigned nine, four, and eight valid alleles to *I. alpestris*, *P. waltl,* and *P. perezi*, respectively. (Figure 1; Additional Figure S1). MHC class II exon 2 length varied between species (239 bp; *I. alpestris*, 270 bp; *P. waltl,* and 252 bp; *P. perezi*) by using the DOC method (Lighten et al., 2014).

### MHC class II exon 2 diversity

We identified 21 unique MHC class II exon 2 alleles among the three species (*I. alpestris* N=9, *P. waltl* N=4; *P. perezi* N=8). Four MHC class II alleles were recovered at relatively high frequency in *I. alpestris* (Ich.alp*01,02,03,04) and a single one was covered at very high frequency in *P. waltl* and *P. perezi*, respectively (Pleu_wa*02 and Pe_pe*01). We did not find any significant differences in allele frequencies between the infection groups (Bd infected, Rv infected, Bd+Rv infected, and non-infected). MHC class II exon 2 presented the lowest gene diversity in *P. waltl*, which ranged from 0.35 to 0.44 compared to *P. perezi* or *I. alpestris* (Table 1). MHC class II exon 2 revealed lower nucleotide diversity (π) but higher nucleotide differences (theta k) in *P. perezi* compared to the other two species (Table 1). Number of private alleles per infection group were present in the three species (Ich_alp*07 for *I. alpestris,*, Pe_pe*07,08 for *P. perezi,* and Pleu_wa*05 for *P. waltl*; See Figure 1 and Additional Table S1). The rest of alleles were shared within the infection group in every species.

We detected two alleles per individual in *P. waltl* while we detected more than two alleles in *I. alpestris* and *P. perezi*, indicating multiple copies in all sequenced MHC class II exon 2 locus in the last two species. Number of valid MHC class II exon 2 alleles per individual varied from one and two copies in *P. waltl*, two to six in *I. alpestris,* and one to four in *P. perezi*. The average number of gene copies per MHC class II exon 2 loci ranged from 1 to 3 (*I. alpestris*; 2.72 ± 0.46 [s.d.], *P. perezi*; 1.23 ± 0.42 [s.d.], *P .waltl*; 1, See Supplementary Figure S3). Traditionally, the number of MHC gene copies in non-model species is estimated by screening MHC variation at the intraspecific level. This is done by detecting the maximum number of putatively functional MHC alleles in each individual and dividing it by two, assuming heterozygosity at each locus (Minias, Palomar, Dudek, & Babik, 2022). This result was verified visually in the haplotype phylogenetic tree provided in order to include this as a variable for the consecutive analyses (Figure 2). The unrooted phylogenetic network showed that the MHC class II exo2 alleles were clustered by species forming monophyletic clades. (See Supplementary Figure S4).

### Supertype and haplotype-supertype diversity

We converted the alleles into six functional supertypes based on physio-chemical binding properties (Figure 2; See (Savage & Zamudio, 2016). After the supertype assignment, all the alleles were monophyletic clustered by species in specific supertypes. However, the branches were widespread over the phylogenetic tree according to its functionality. The MHC supertype S2 and S4 were assigned to *P. waltl* and *P. perezi*, respectively while S1, S3, S5, and S6 supertypes were assigned to *I. alpestris* (Figure 2). We identify seven MHC haplotype-supertypes in *Ich.alpestris*, of which one was found at high frequency (Ich_alp*2_1_0_0; >50%, See Supplementary Figure S6).

### MHC copy number variation and infection load among study species

Our results show that *Bd* infection values after standardization remain relatively low in all species. *Bd* infection load was significantly different between species (X^2^=18.2580, Df=2, P=0.0001) and infection groups (*Bd*_infected and *Bd*+*Rv*_infected; X^2^= 5.7532, Df=1, P= 0.016, See Supplementary Figure S8). For the *Rv* infection, we found significant differences in *Rv* infection load between species (X^2^=50.0059, Df= 2, P> 0.0001) but not between infection groups. *Rv* infection was present in *I. alpestris* but at very low intensities in comparison to *P. perezi* and *P. waltl* (*P. perezi*; SE=1.5336, Z=6.179, P<0.0001, *P. waltl*; SE=1.4798, Z=6.288, P<0.0001; Supplementary Figure S8). Regarding the number of loci, we found that *Bd* infection load was significantly different depending on the number of loci that the individuals were carrying (X^2^=13.25, Df=2, P= 0.0013), but not for *Rv-* infection (X^2^=4.25, Df=2, P= 0.119) or between infection groups (X^2^=13.25, Df=2, P= 0.0013; See Figure 3).

### The effect of specific MHC class II exon 2 configuration on infection

GLM analyses of infection status (1; Infected, 0; Non-infected) found that individuals carrying the allele Ich_alp*04 had a significant effect on general infection (LR X^2^=8.76, Df=1, P=0.003), *Rv-* Infection (LR X^2^= 5.9381, Df=1, P=0.01), and co-infection (LR X^2^= 8.7599, Df=1, P=0.004) in *I. alpestris*. For Bd infection, this effect was nearly significant (LR X^2^= 3.1965, Df=1, P=0.07). The general effect of MHC alleles on infection may be likely positive when individuals are infected with *Bd*, *Rv* or co-infected (see the coefficient regression plot; Supplementary Figure S9). We did not find any association between Pe_pe* alleles over general or specific infection/co-infection in *P. perezi* species.

Additionally, GLMM analyses on infection status showed that S1 (Supertype 1) only present in *I. alpestris* was significantly associated with infection (X^2^= 4.0856, Df=1, P=0.043: Supplementary Figure S10). We found that the effect of S1 on infection is directly related to the presence of Ich_alp*04 because it belongs to the same functional supertype group. Although we did not find any association between haplotype-supertypes and infection, descriptive plots show that two *I. alpestris* individuals that carry the haplotype supertype Ich.alp*2_1_0_0*P and Ich.alp*2_1_1_1*C does not present any infection (Supplementary Figure S11). Also, Ich.alp*2_2_0_2*E haplotype supertype is only present in infected individuals (Supplementary Figure S11).

## DISCUSSION

Our study provides the first characterization of MHC class II diversity in three amphibian species in a co-infection scenario, while previous MHC-pathogen studies on amphibians have considered only one pathogen. Both, the immunogenetic optimality hypothesis and associations between specific pathogens and particular MHC alleles were evidenced. The associations between MHC allele (Ich.alp*04) and supertype “S1” on infection status for *Ranavirus* and co-infection in *I. alpestris* suggest a substantial role for either rare-allele advantage or positive selection and fluctuating selection for that species (Spurgin & Richardson, 2010). Therefore, monitoring the frequencies of theses alleles in the population might give us more evidence to support the selective mechanisms acting on MHC class II. The significant association between MHC genotypes and *Bd* and *Rv* infection status observed in *I.alpestris* individuals provides evidence that MHC class II variants are indeed contributing to antiviral and antifungal immunity in amphibian populations (Savage, Muletz-Wolz, Campbell Grant, Fleischer, & Mulder, 2019). However both negative frequency-dependent selection and fluctuating selection lead to associations between pathogens and specific alleles, making hard to disentangle between these two different mechanisms by observational studies alone. The associations between MHC allele (Ich.alp*04) and supertype “S1’’ on infection prevalence for *Rv* could be explained by fluctuating selection, either temporally or spatially, however in our study we did not analyzed temporal and spatial heterogeneity, which could be very informative because annual variation in pathogen prevalence may be related to both climate and host factors (Savage & Zamudio, 2011). Climatic conditions are known to affect pathogen survival in the environment and may influence host transmission (Metcalf et al., 2017). Alternatively, amphibian populations are characterized by structuring geographically (Alex Smith & M. Green, 2005; Watts et al., 2015), how populations are spatially aggregated and connected can have a profound impact on the population dynamics and evolutionary landscapes of species. Including a spatial and temporal perspective on the dynamics of the host and pathogen populations would help to interpret the results.

MHC has been characterized in a great variety of taxa (fish (Edholm, Pasquier, Wiegertjes, & Boudinot, 2022); primate, (Heijmans, de Groot, & Bontrop, 2020); ungulates, (Ivy-Israel, Moore, Schwartz, & Ditchkoff, 2020); and reptiles, (Reed & Settlage, 2021), including many amphibian species (Cortázar-Chinarro et al., 2017); (Kiemnec-Tyburczy, Richmond, Savage, Lips, & Zamudio, 2012); (Palomar et al., 2021); (Talarico, Babik, Marta, Pietrocini, & Mattoccia, 2019). (Babik, Pabijan, & Radwan, 2008) have previously characterized MHC II in *I.alpestris* from the South of Poland, showing relatively high genetic variation and a maximum of nine alleles per individual, implying the existence of at least five loci in this species in Eastern Europe. We have only found a maximum of six alleles per individual coinciding with the existence of at least three loci in the same species. Whether this difference comes from the number of loci or from the sharing of alleles between MHC copies in Spanish population is unknown. Further immunogenetic studies are needed to characterize MHC genetic diversity in *I.alpestris* throughout its geographical distribution. Alternatively, this study characterized for the first time MHC II in *P. perezi* and *P. waltl*. MHC II genetic diversity is high for *P. perezi* but quite low for *P. waltl*. In this last species, we only found four different MHC alleles (Pleu_wa*01,02,05,07) and two out of four were presented at very low frequency (Pleu_wa*05,07). The recent publication of the P. waltl genome (Brown et al., 2022) has revealed the presence of at least one MHC II gene in this species. This finding suggests that *P. waltl* may have lower genetic diversity in this region compared to other species that carry multiple MHC II genes. Notably, our analysis of 57 individuals revealed only four alleles for this gene, which is typically associated with high levels of diversity. This paucity of variation may be a consequence of mass mortalities attributed to Bd or Rv, which could have selectively favored a few resistant alleles. Further investigations comparing populations affected by emerging diseases with those that are not could provide valuable insights into this matter. II

Previous studies investigated the associations between specific MHC alleles or supertypes and pathogens in amphibians (Bataille et al., 2015; Cortazar-Chinarro et al., 2022; Savage & Zamudio, 2011). Howerver, the nature of these associations and potential contribution in amphibian anti-viral/fungal immunity remain poorly understood due to the scarce number of experimental infection studies in amphibian species (See e.g (Bataille et al., 2015; Cortazar-Chinarro et al., 2022), as an example). A clear step forward might be to investigate whether variation in functional immunity correlates to *Bd/Rv* infection load and prevalence across different anuran taxa, both within and between species, to ultimately investigate the potential role of specific functionally expressed MHC genotype on infection (Savage et al., 2019). For instance, we recommend a multi-level studies implementation: 1) immune gene diversity, 2) functional diversity and 3) experimental infection studies in single and co-infected species to ultimately study whether specific MHC allele configurations are positively/negatively associated with mortality, survival or individual fitness.

Our results also showed that many individuals of *P. perezi* and *I. alpestris* carried multiple copies of MHC gene, suggesting single or multiple gene duplication events over the amphibian species evolutionary history. We also demostrate that the number of loci that an individual carries is significantly associated with infection intensity although we did not find a linear relationship. Previous studies have shown that individuals with a great number of MHC alleles display lower parasite loads matching the heterozygote advange hypothesis (Meyer-Lucht & Sommer, 2009). We detected a higher infection intensity for individuals that carry one single locus than for individuals that carry three or more, suggesting that having one MHC copy or more than three might potentially have a negative effect on the host (e.g a decrease in survival) when confronting infections. Nevertheless, experimental infection studies could provide further validation for this assertion. We did not find a negative linear relationship between MHC diversity and status or intensity of infection. However, we found that individuals that carried intermediate number of MHC loci had a lower infection intensity compared to the ones carrying a single MHC locus, three or more MHC loci, in concordance with the immunogenetic optimality hypothesis. This hypothesis argues that, at the individual level, it is not maximized but rather optimized heterozygosity that provides maximum fitness benefits (Nowak et al., 1992) and has been demonstrated in other cases (Kloch et al., 2010; Madsen & Ujvari, 2006; Wegner et al., 2003; Wegner, Kalbe, Milinski, & Reusch, 2008). On the contrary, other studies have not found such a link between lower infection load and intermediate number of MHC alleles (Garamszegi et al., 2015; Radwan et al., 2012; Stervander, Dierickx, Thorley, Brooke, & Westerdahl, 2020), or even argue that might be context dependent (Råberg et al., 2022). In any case, it seems that Bd and Rv infections could select MHC alleles in amphibians promoting intermediate heterozygosity.

While the immunogenetic optimality hypothesis matched with our results for *Bd-infection*, we did not find any significant relationship between number of MHC loci and infection load in *Rv-infected* individuals or co-infected individuals. Despite examining co-infection in three different species we found no support for the MHC heterozygote advantage in relation to co-infection, which could be explained because in a multiple pathogen context MHC alleles conferring resistance to one pathogen can increase susceptibility to another (Penn & Potts, 1999). Among previous work done with chytridiomycosis, there is a great disparity of results, some have found evidence that MHC heterozygosity contributes to to chytridiomycosis resistance (Savage & Zamudio, 2011), while others did not observe any evidence of heterozygote advantage (Bataille et al., 2015) or even found an homozygous advantage (Savage, Mulder, Torres, & Wells, 2018). However, we are not aware that other studies have tested this hypothesis in amphibians in a co-infection scenario, in which individuals must be able to resist two different pathogens. Further studies of wild populations exposed to multiple pathogens are needed to better understand the evolution of the amphibian immunogenetic system. Work carried out with other organizations has provided evidence of the advantage of heterozygotes to co-infections with multiple pathogens (McClelland, Penn, & Potts, 2003; Sin et al., 2014).

We speculated that the lack of relationship between the number of MHC loci and *Rv* infection load might be due to different reasons. We should consider the differences in the evolutionary history of the parasites (*Bd* and *Rv*) in the Iberian Peninsula. Amphibian declines due to chytrid fungus have been reported since mid 1990’s in the Iberian Peninsula (Bosch & Martínez-Solano, 2003, 2006), while *Rv* mass mortalities were recorded for the first time in 2005 (Price et al., 2014). Despite the enormous coexistence for years between both pathogens, evolutionary life history of *Bd* and *Ranavirus* might suggest a different immune response in the host (Fu & Waldman, 2017; Warne et al., 2016). Second, the establishment/adaptation of parasites on a specific habitat might indirectly contribute to the evolution of immune genes and it might be quite different for *Bd* and *Ranavirus*. For instance, (Hoverman, Mihaljevic, Richgels, Kerby, & Johnson, 2012) speculated that the time that an amphibian spend in the water might be related to host susceptibility to multi-host pathogens and therefore it might be indirectly related to the host immune response. Additionally, sticklebacks showed local adaptation of MHC genotypes to population-specific parasites, independently of the genetic background (Eizaguirre, Lenz, Kalbe, & Milinski, 2012). To date, the association between MHC copy number variation and infection intensity and the implications of the complex host-parasite dynamics in nature remains unresolved.

Our study illustrates the importance of looking beyond single snapshot MHC/pathogen association in wildlife amphibian immunogenetic studies. In the present case, we did not find any evidence of the heterozygote advantage hypothesis in the three species in relation to co-infection, while we found an association between a specific allele number and the infection intensity, supporting the immunogenetic optimal hypothesis. Further NGS-based genotyping methods that allow testing for the optimal hypothesis are in urgent need in order to address this open gap in a variety or other, non-model organisms and specially in amphibian species.

## AUTHOR CONTRIBUTIONS

CM: did all wet lab work, analyses, conceptualization, funding and wrote the first draft of the manuscript; ALB: wrote the first draft of the manuscript, PH; analyzes support, GP; conceptualization of study and primer design support, BJ field work-field work funding and conceptualization of the study, HW; host institution representative mentor and funding. All authors contributed to reviewing and editing

## AKNOWLEGDMENTS

The authors would like to acknowledge Alba Cortazar Chinarro for the amphibian water paint illustrations present in the manuscript. This study was supported by the Swedish Research Council (M.C.-C.International mobility grant; 2019-06352).

## DATA AVAILABILITY STATEMENT

All data have been made available in: pending upload

