## Supplementary material for "The Major Histocompatibility Complex modulates *Batrachochytrium dendrobatidis* and *Ranavirus* infections in three amphibian species": suplementary Material

^6^Departamento de Biomedicina y Biotecnologia, Universidad de Alcala de Henares, Pza. San Diego s/n, 28801 – Alcalá de Henares (Madrid), Spain.

^7^Instituto Mixto de Investigacion en Biodiversidad, Universidad de Oviedo, Campus de Mieres. Edificio de Investigacion- Planta 5, c/ Gonzalo Guiterrez Quir´s s/n, 33600 Mieres, Asturias.

**ADDITIONAL FIGURES AND TABLES**

**TABLES**

**Table S1.** Miseq run summary for the two independent Miseq runs. (N) is defined as the total number of samples included in the study. The percentage (%) of duplicated is directly related to the number of replicates out of the total number of samples in the study. The 3% of the average of reads calculated from the average number of reads per sample.

| **samples** |  |
| --- | --- |
| (N) *I. alpestri*s | 69 |
| (N) *P. perezi* | 51 |
| (N) *P. waltl* | 57 |
| **Miseq Run 1 *I. alpestris*** |  |
| Total Number of reads | 2,100,296 |
| Total (N) | 90 |
| Average number of reads per sample | 23,336.62 |
| % of duplicates | 31.88 % |
| 3% of the average number of reads | 700.09 |
| Final retained samples | 68 |
| **Miseq Run 2 *P. perezi* *P. waltl*** |  |
| ***P. perezi*** |  |
| Total Number of reads | 3,132,387 |
| Total (N) | 108 |
| Average number of reads per sample | 29.003,58 |
| % of duplicates | 100 |
| 3% of the average number of reads | 870 |
| Final retained samples | 46 |
| ***P. waltl*** |  |
| Total Number of reads | 443,215 |
| Total (N) | 171 |
| Average number of reads per sample | 2591.90 |
| % of duplicates | 100 |
| 3% of the average number of reads | 77.75 |
| Final retained samples | 51 |

**FIGURES**

**Figure S1**. Allele color scheme for the three different species*: I. alpestris, P. perezi, and P. waltl*

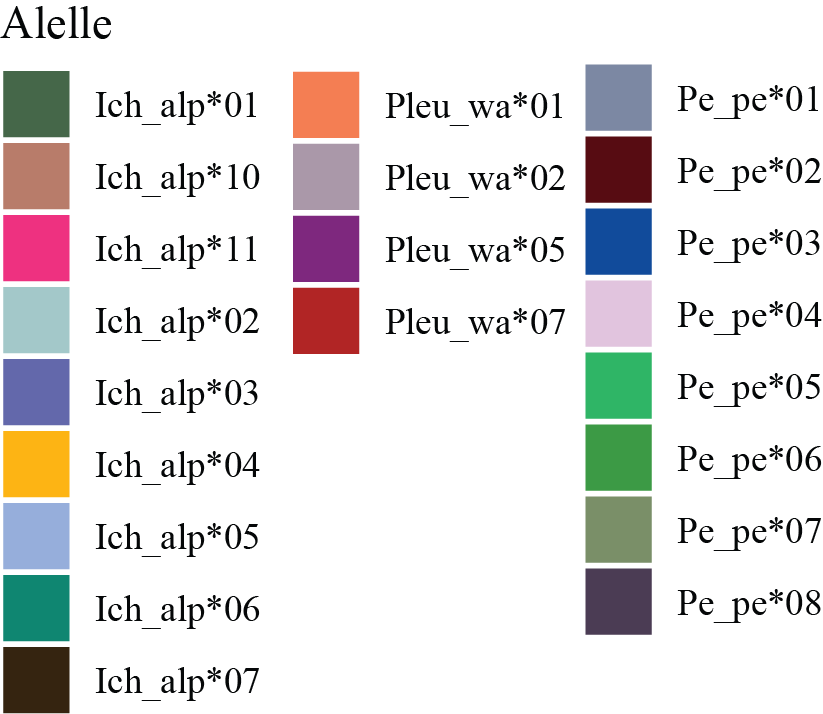

**Figure S2**. Allele frequency per infection group: A) *I. alpestris*, B) *P. waltl* and C) *P. perezi*

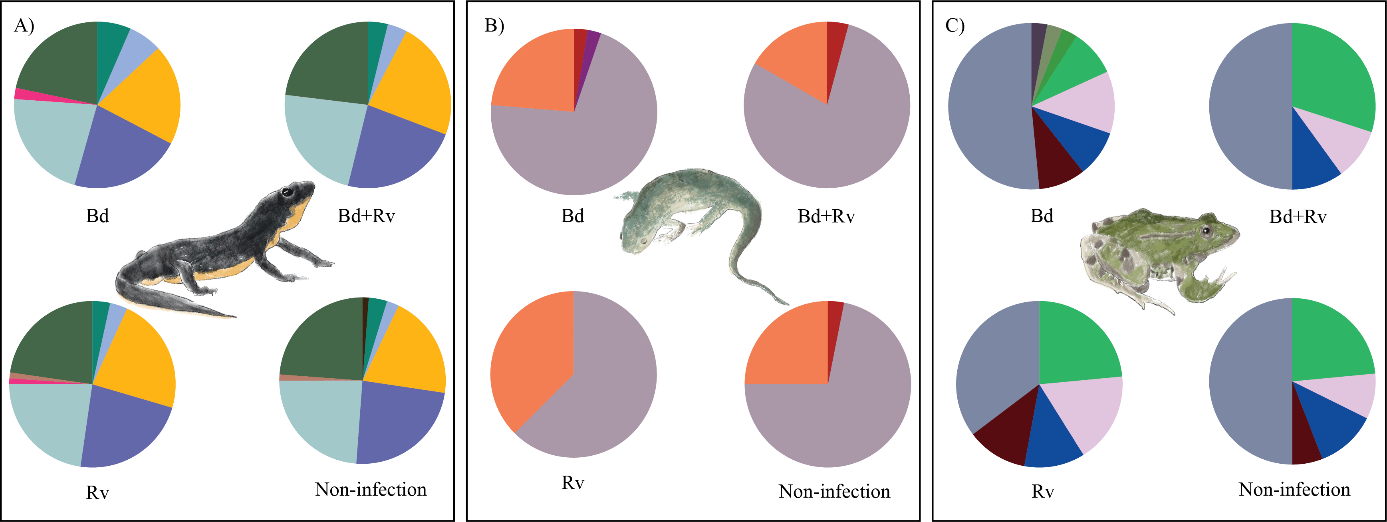

**Figure S3**. Molecular phylogram of nucleotide sequences of MHC class II exon 2 reconstructed with neighbor joining methods for the three species A) *I. alpestris*, B) *P. waltl* and C) *P. perezi*
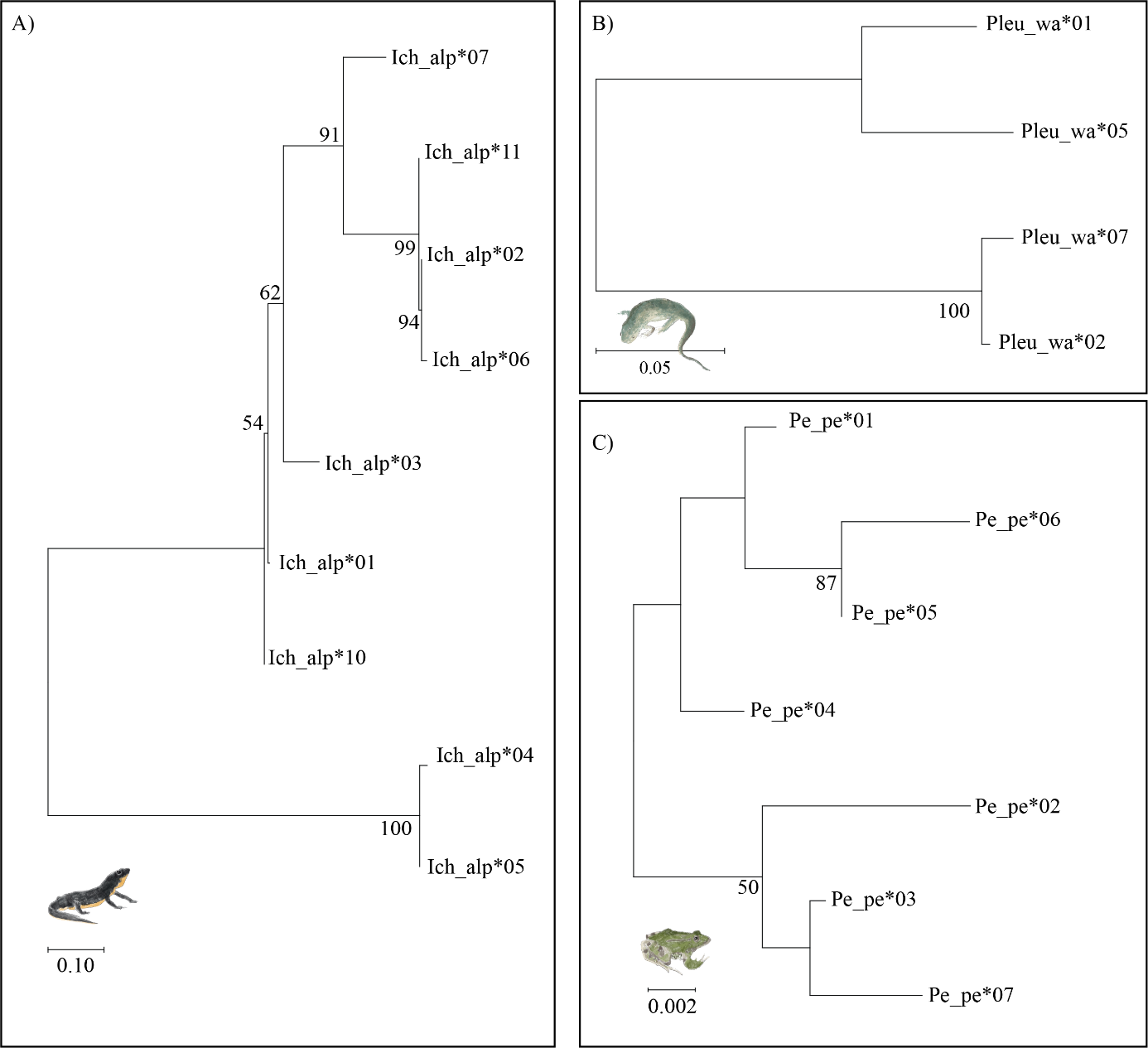
. Bootstrap values from 1000 replicates greater than 50% are indicated on branches. The valid alleles were named following the nomenclature by Klein (1975) for MHC loci: a four-digit abbreviation of the species name followed by species_gene*numeration, e.g .Ich_alp*01.

**Figure S4**. Unrooted phylogenetic network to illustrate the phylogenetic relationship between the amphibian species. We included a *Rana arvalis* sequence from Cortazar-Chinarro et al. 2017, as an outgroup.

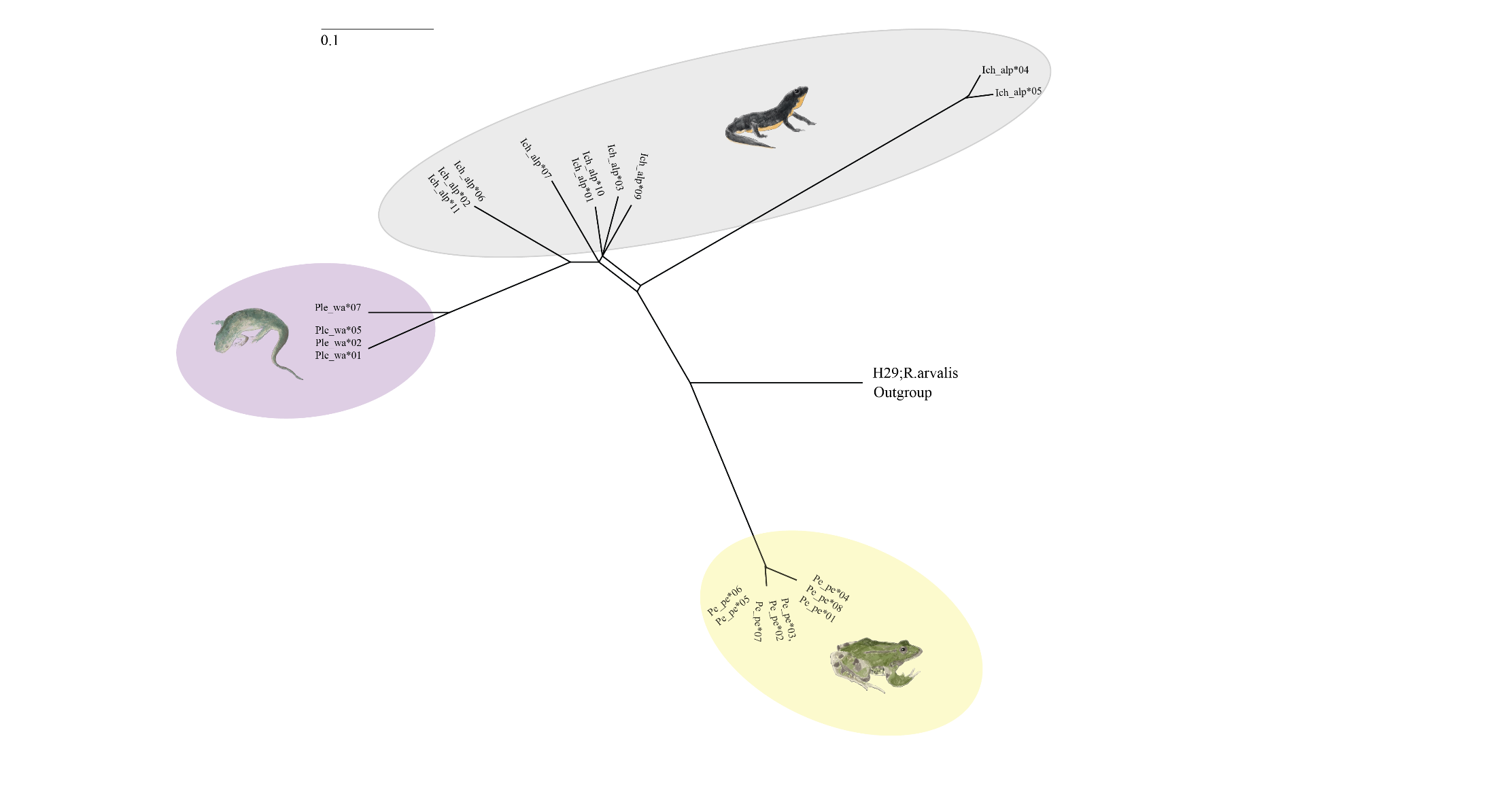

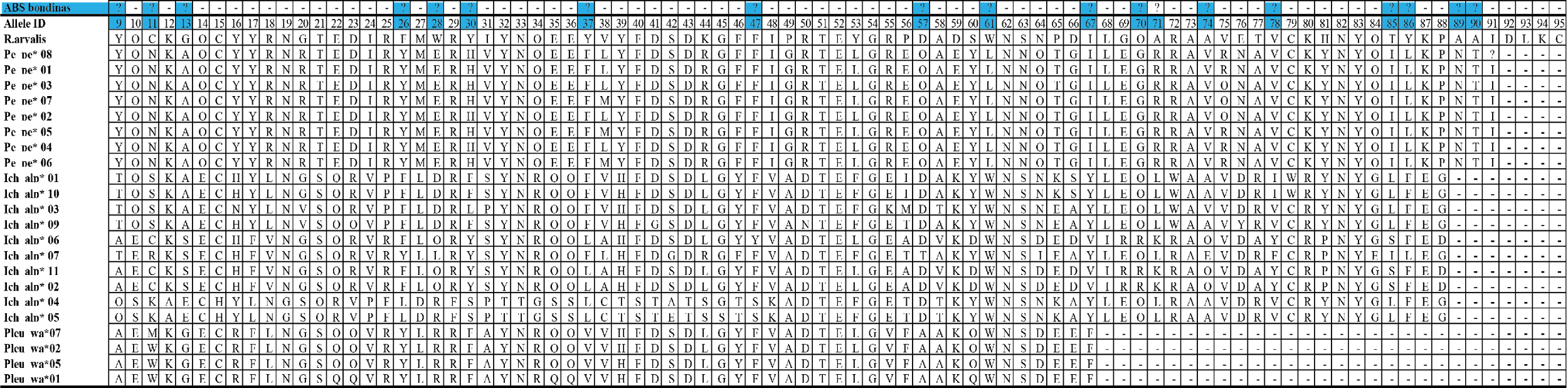
**Figure S5.** Alignment of amino acid sequences. PBR positions from Bondinas et al. 2007 are marked with (?) in blue

**Figure S6**. Supertype_haplotype diversity in *I. alpestris*

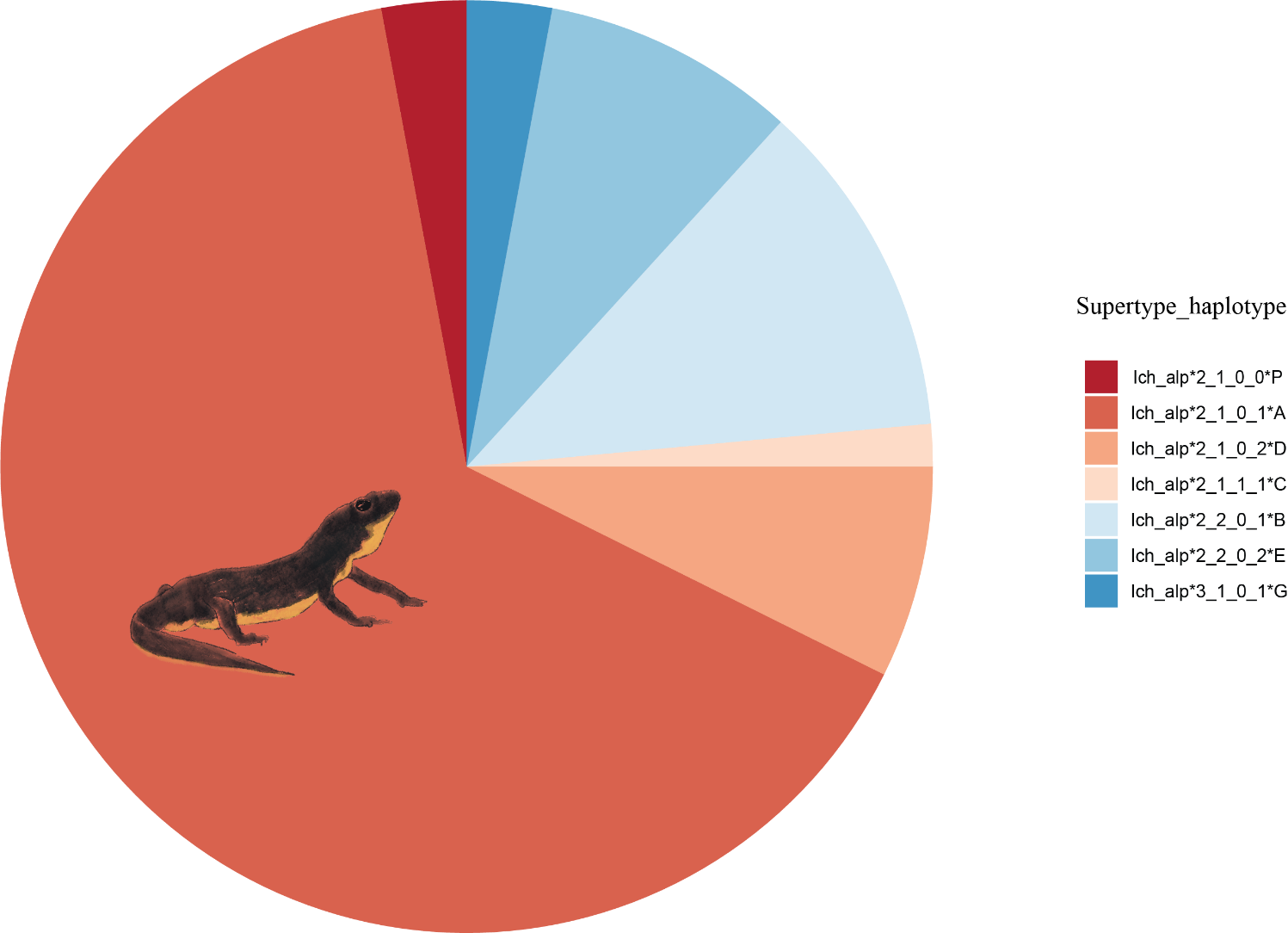

**Figure S7.** Descriptive bar plot comparing A) 1- number of individuals in each infection category by location, 2- number of individuals in each infection category by number of loci, B) number of individuals count carrying one, two or three MHC class II exon 2 loci in three different species: *I. alpestris, P. perezi,* and *P. waltl*. Note that the number of loci has been calculated as half of the number of alleles.

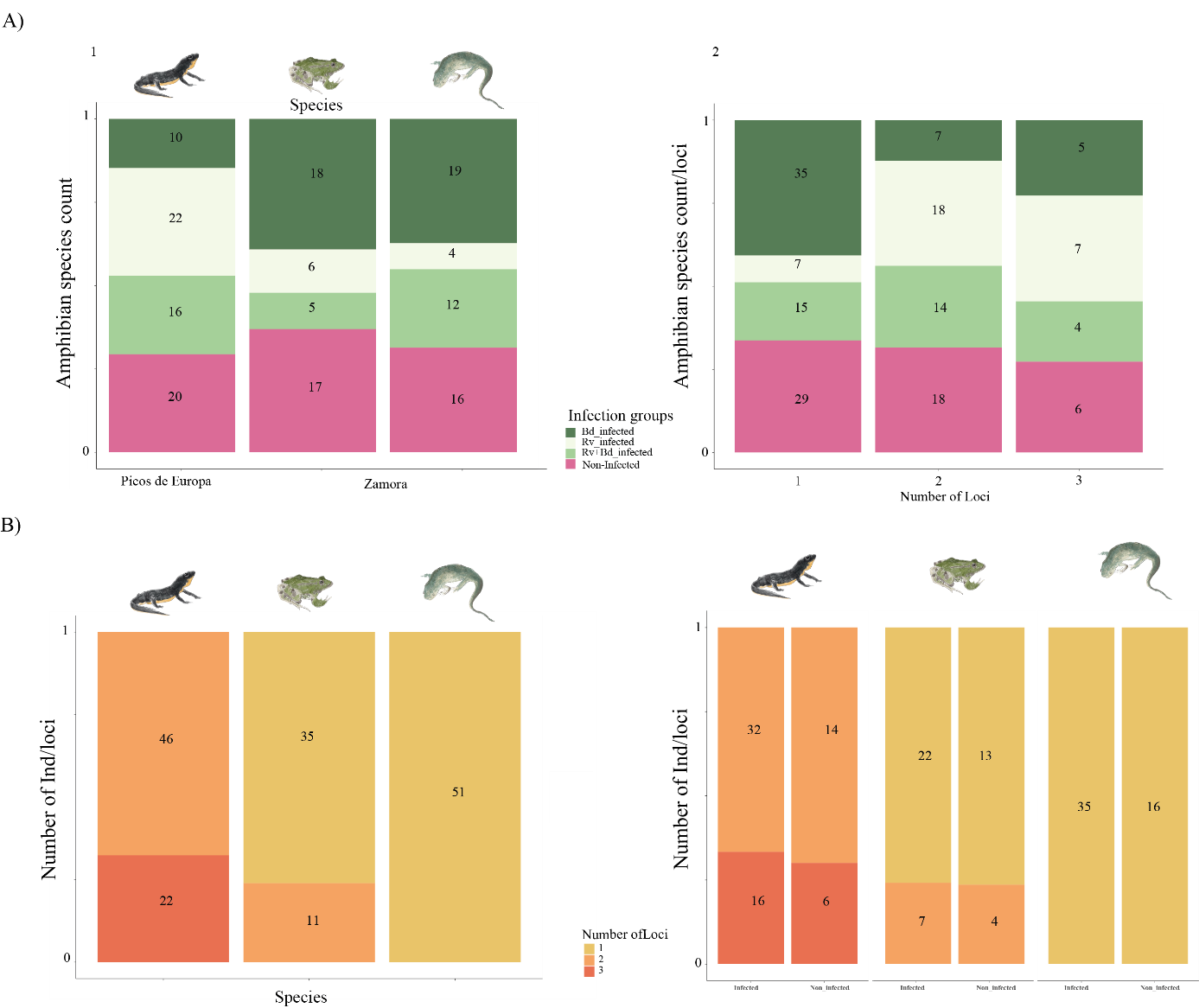

**Figure S8.** Infection load comparations a) for *Bd* infection including *Bd* infected individuals and co-infected individuals and b) for *Rv* infection including *Rv* infected and co-infected indiduals in the three species included in the study: *I.alpestris, P. perezi* and *P. waltl*. Significant p-values in infection load between species are marked with * based on the GLMM analyses performed with CRAN package (glmmTMB) developed by Magnusson et al. (2017).

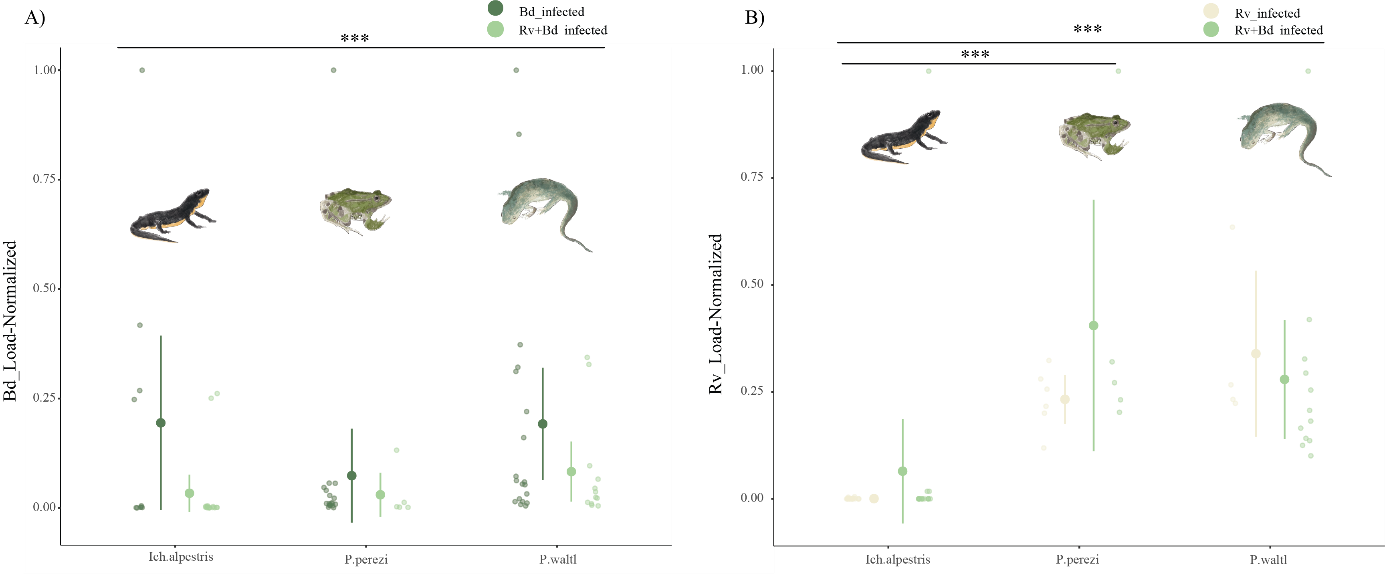

**Figure S9.** Regression coefficient plots representing the main effect of carrying a specific MHC class II exon 2 allele over infection. Models included in the figure are based on presence/absence (1, 0) and are: Infection/non-infection, *Bd*-infected/non infected, *Rv*-infected/non infected, and co-infection (*Bd*+*Rv*)/not coinfection. Significant alleles are represented with a *

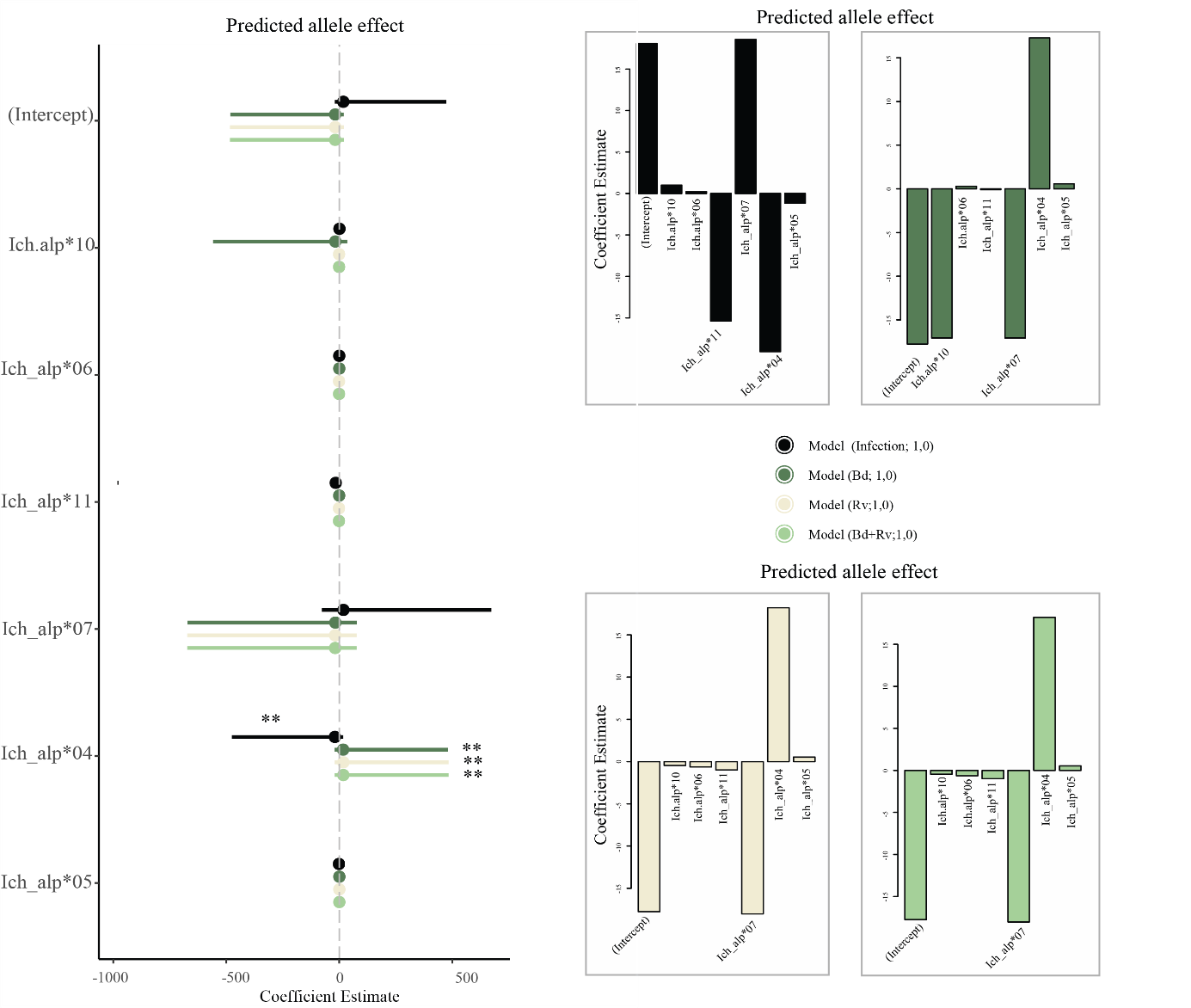

**Figure S10.** Regression coefficient plot representing the main effect of carrying a specific MHC class II exon 2 supertype in *I.alpestris* over infection (infected –non-infected individuals; 1, 0). Significant supertypes are marked with an (*).

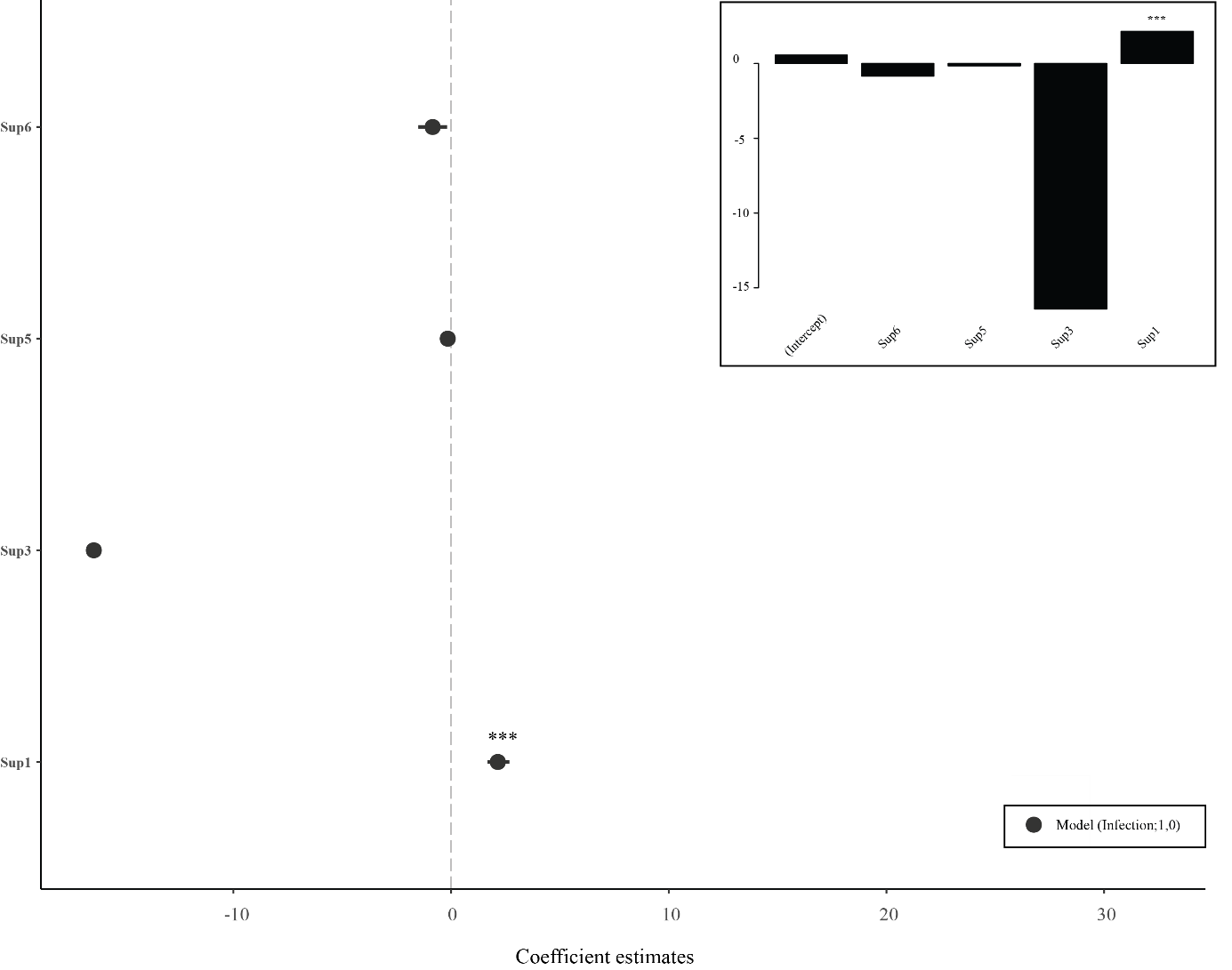

**Figure S11.** Regression coefficient plot representing the main effect of carrying a specific MHC class II exon 2 supertype- haplotype over infection (infected –non infected individuals; 1,0).

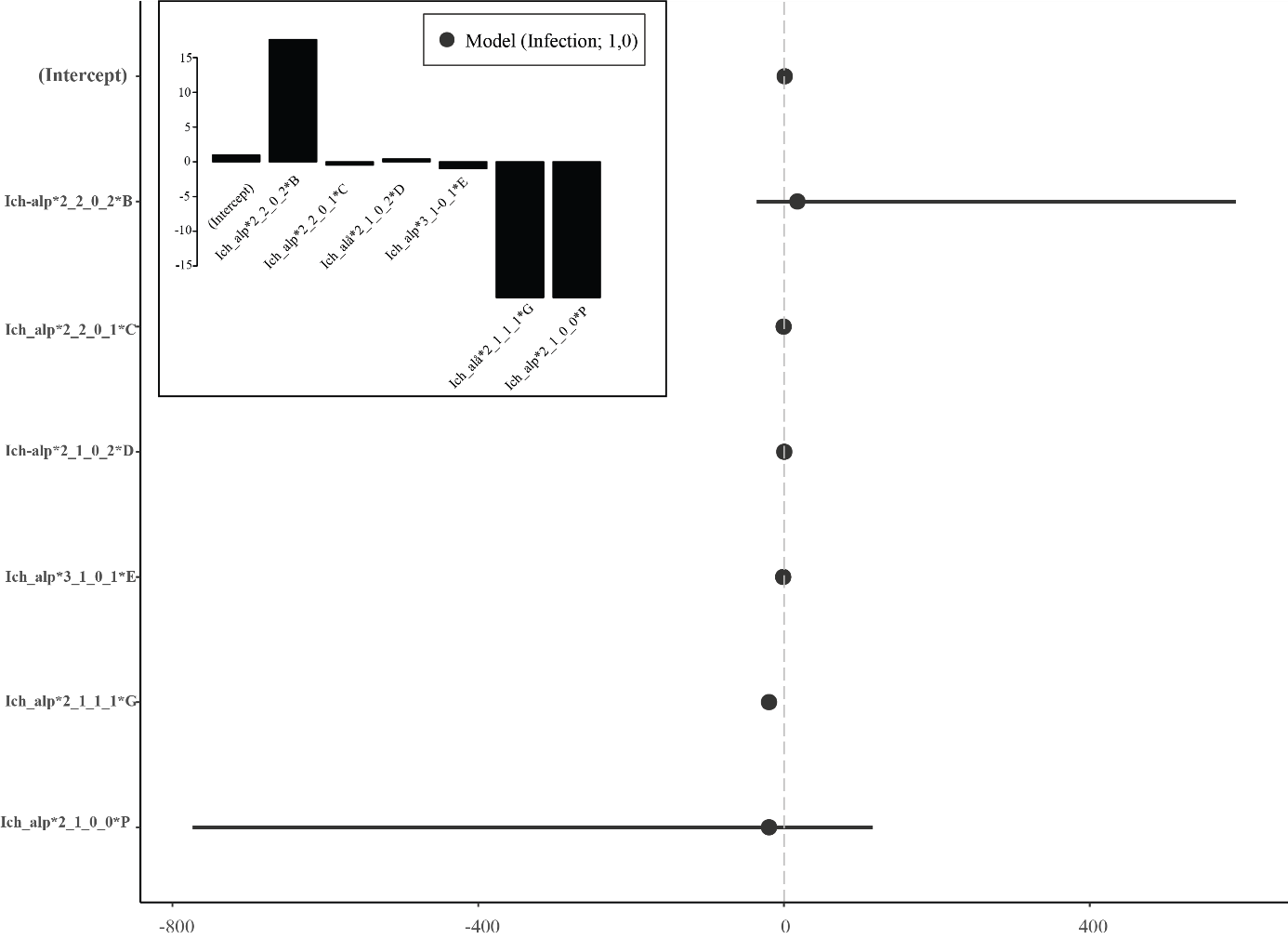

**Figure S12.** Descriptive bar plot representing A) number of individuals carrying an specific MHC class II exon 2 supertype-haplotype considering four infection categories: *Bd*-infected, *Rv*-infected, *Rv*+*Bd* infected and uninfected, B) number of individuals carrying and specific MHC class II exon 2 supertype-haplotype over infection (1,0).

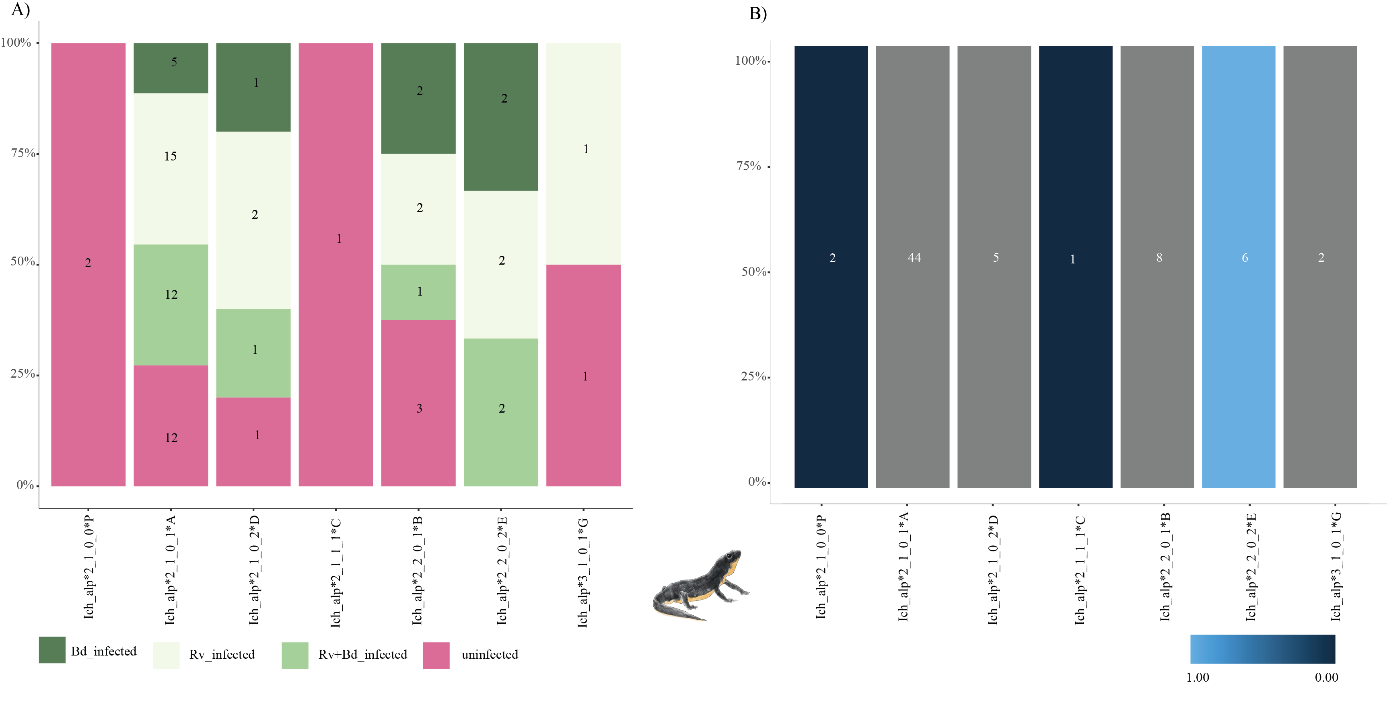
